# Functional characterization of *Polr3a* hypomyelinating leukodystrophy mutations in the *S. cerevisiae* homolog, *RPC160*

**DOI:** 10.1101/2020.06.30.180125

**Authors:** Robyn D. Moir, Christian Lavados, JaeHoon Lee, Ian M. Willis

## Abstract

Mutations in RNA polymerase III (Pol III) cause hypomeylinating leukodystrophy (HLD) and neurodegeneration in humans. POLR3A and POLR3B, the two largest Pol III subunits, together form the catalytic center and carry the majority of disease alleles. Disease-causing mutations include invariant and highly conserved residues that are predicted to negatively affect Pol III activity and decrease transcriptional output. A subset of HLD missense mutations in POLR3A cluster in the pore region that provides nucleotide access to the Pol III active site. These mutations were engineered at the corresponding positions in the *Saccharomyces cerevisiae* homolog, Rpc160, to evaluate their functional deficits. None of the mutations caused a growth or transcription phenotype in yeast. Each mutation was combined with a frequently occurring pore mutation, POLR3A G672E, which was also wild-type for growth and transcription. The double mutants showed a spectrum of phenotypes from wild-type to lethal, with only the least fit combinations showing an effect on Pol III transcription. In one slow-growing temperature-sensitive mutant the steady-state level of tRNAs was unaffected, however global tRNA synthesis was compromised, as was the synthesis of *RPR1* and *SNR52* RNAs. Affinity-purified mutant Pol III was broadly defective in both factor-independent and factor-dependent transcription *in vitro* across genes that represent the yeast Pol III transcriptome. Thus, the robustness of yeast to Pol III leukodystrophy mutations in *RPC160* can be overcome by a combinatorial strategy.

## 1. Introduction

Transcription by RNA polymerase III (Pol III) is an essential process that generates stable non-coding RNAs involved in protein synthesis, RNA processing, protein secretion and other processes (Dieci et al., 2007). The majority of Pol III-transcribed genes encode tRNAs, the amino acid adaptors that decode mRNA sequences during protein synthesis. The Pol III transcription machinery comprises essential multi-subunit complexes that include the 17-subunit polymerase and a three-subunit initiation factor, TFIIIB. Biallelic missense mutations in either *Polr3a* or *Polr3b* (which encode the two largest subunits of Pol III), *Polr3k*, or *Polr1c* (a shared subunit of Pols I and III) cause childhood-onset hypomyelinating leukodystrophy (HLD) (Wolf et al., 2014; Gutierrez et al., 2015; Thiffault et al., 2015; Dorboz et al., 2018). Pol III HLD mutations are anticipated to alter Pol III transcription throughout the body but the primary phenotypic effect is on the central nervous system. The major feature of this early-onset autosomal recessive disease is hypomyelination of cerebral white matter. Missense mutations in the TFIIIB subunits, BRF1 and BDP1, have also been linked to neurological disease, causing cerebellar hypoplasia and cognitive dysfunction (Borck et al., 2015) and early-onset severe hearing loss, respectively (Girotto et al., 2013; Girotto et al., 2014). Disease-causing mutations in Pol III and initiation factor subunits include invariant and highly conserved residues that are predicted to negatively affect Pol III activity and decrease transcriptional output (Trinh et al., 2006; Bernard et al., 2011).

The mechanisms of pathogenesis of Pol III HLD disease mutations are unknown, but decreased Pol III transcription is anticipated to alter the protein synthetic capacity of cells. Studies of the Pol III transcriptome in HLD patient-derived samples report varied dysregulation of Pol III gene expression and gene-specific sensitivity to POLR3 mutations (Dorboz et al., 2018; Choquet et al., 2019a). Eight patient-derived fibroblast lines, representing homozygous and compound heterozygous *Polr3a* HLD genotypes showed a common decrease in BC200 RNA, a brain-specific RNA linked to repression of translation (Choquet et al., 2019a). Two fibroblast lines derived from homozygous mutant *Polr3k* HLD patient samples showed lower levels of 5S rRNA, 7SL RNA and tRNAiMet levels (Dorboz et al., 2018). Splicing mutations in *Polr3a* are linked to a striatal variant of POLR3A-related disease that is clinically distinct from HLD: An analysis of blood samples from three such patients showed a reduction across all tRNAs, negative effects on the steady-state abundance of a subset of other Pol III transcripts, and elevated 5S rRNA, and RppH1 and 7SK RNAs (Azmanov et al., 2016). The Pol II transcriptome was also altered in these samples, with a reduction of chaperone protein expression and enrichment in snoRNA maturation proteins.

Experimental models of Pol III HLD report varied defects in Pol III gene expression. Expression of a POLR3A M852V disease mutation in a human cell line resulted in a global decrease in precursor tRNA levels and lower steady state levels of BC200 RNA and 7SL RNA. Other Pol III-transcribed genes were unaffected, as were the levels of mature tRNAs (Choquet et al., 2019a). Expression of the M852V mutation in an oligodendroglial cell line, that can be induced to differentiate from preoligodendocyte-like cells to cells that express myelin basic protein (MBP), resulted in decreased BC200 RNA and MBP mRNA levels. While the proteome of these cells showed both positive and negative effects on a limited number of proteins, there was no global effect on protein abundance (Choquet et al., 2019a). Expression of two POLR3A disease mutations in the *S*.*pombe* homolog of POLR3A resulted in a decrease in several precursor tRNAs, elevated m^2^_2_G26 modification of a subset of tRNAs and enhanced tRNA decoding of a reporter gene, thought to be due to the increased stability of the mature tRNA (Arimbasseri et al., 2015). This change in tRNA modification was induced in human cells by a nutrient stress-mediated shutdown of transcription, indicating that the tRNA modification enzyme Trm1 is broadly limiting for function. These reports indicate that Pol III disease mutations in patient-derived samples and in experimental models compromise the synthesis of a subset of the Pol III transcriptome and can have specific effects on protein output.

Pol III HLD remains poorly characterized at the cellular and molecular level. Mouse models of Pol III-related HLD have proven difficult to generate. Mice homozygous for the most frequent Pol III-associated HLD missense mutation, G672E in POLR3A, did not recapitulate human disease phenotypes either as pups or at advanced age. Moreover, no defect in Pol III transcription was detected in the mice or a knock-in human cell culture model (Choquet et al., 2017). Mice in which the G672E mutation was combined with a *Polr3a* null allele also did not exhibit neurological abnormalities or transcription phenotypes (Choquet et al., 2017). In contrast, a POLR3B R103H mutation that impairs the assembly of Pol III when ectopically expressed in human cell lines, was viable as a heterozygote knock-in in the mouse but embryonically lethal as a homozygote (Choquet et al., 2019b). A double-mutant mouse, with a *Polr3a*^*G672E/null*^ *Polr3b*^*+/R103*^ genotype, also did not exhibit neurological or transcription phenotypes (Choquet et al., 2019b). Together, these phenotypes suggest that genetic robustness limits the effect of Pol III mutations and/or deficits in Pol III function in the mouse. Alternatively, genetic modifiers and/or environmental stressors in human development may contribute to HLD disease presentation and depth of phenotype (Choquet et al., 2017).

The phenotypic disparity between the human disease and mouse models of Pol III-associated HLD led us to screen a panel of POLR3A HLD mutations, which map to the pore region where multiple mutations are clustered, and assess their functional deficits in yeast Pol III transcription. Each POLR3A HLD-associated mutation was engineered individually or in combination with the frequently occurring pore mutation (G672E in humans) into the *S*.*cerevisiae* homolog, *RPC160* (G686E, yeast amino acid numbering), and assayed for growth and transcription phenotypes. All HLD-associated single mutations were indistinguishable from wild-type in growth and transcription and all had wild-type steady state levels of 5S rRNA and mature tRNAs. Double mutations showed a spectrum of growth phenotypes as well as lethality. Characterization of one double mutant, with Y685K and G686E mutations, showed defective tRNA synthesis and low steady-state levels of precursor and mature forms of *RPR1* and *SNR52* RNAs *in vivo. In vitro* analysis of affinity-purified mutant Pol III showed it to be defective in factor-independent transcription and in factor-dependent transcription across genes that represent the yeast Pol III transcriptome.

## 2. Materials and Methods

### 2.1 Mutagenesis and subcloning

Rpc160 containing a C-terminal triple hemagglutinin (HA) tag was cloned from a genomically-tagged strain (Lee et al., 2015) into pRS315 and pRS316. Mutations in the pRS315-Rpc160 plasmid were generated using the Gibson Assembly Cloning Kit (NEB) and clones were confirmed by sequencing the entire coding region. Chromosomal deletion of *RPC160* in strain W303 pRS316-*RPC160* used *pFA6a-HphNT1* and standard PCR-based methodology (Janke et al., 2004). *RPC160* mutants were subcloned into pRS405, resequenced and the plasmid linearized with *Ppu*M1 for integration at the *LEU2* locus. Colonies resistant to 5-fluoroorotic acid (FOA) were identified and integration of the mutant allele was confirmed by PCR and sequencing. For multicopy expression of *RPR1*, 160 base pairs upstream and 199 base pairs downstream of the *RPR1* coding region was cloned into pRS423. The wild-type *SSD1* gene in pRS316 included 400 base pairs upstream and downstream of the coding sequence and was obtained from Ted Powers (pPL092).

### 2.2 RNA analysis

Yeast strains, listed in Table S1, were derived from W303 and grown in YPD or synthetic complete drop-out media as required. Yeast spot images were generated using 10-fold serial dilutions of freshly grown cultures diluted to OD_600_ 0.5. Doubling times of yeast strains were determined by real time monitoring in a Bioscreen C instrument and represent the average of technical triplicates and multiple biological replicates for each strain (Moir et al., 2012). Cultures for RNA analysis were grown to OD_600_ 0.4-0.5 at 30° before RNA extraction. Hot-phenol extracted total RNA (10 μg/lane) was processed, hybridized, imaged and quantified as described in (Li et al., 2000a). Probe signal intensities were normalized to that of SNR17a/b RNA (U3). Oligo probe sequences are listed in Table S2. For uracil pulse-labeling, cells (10 OD_600_) grown in synthetic complete media without uracil, were washed and resuspended in 900 μl warm media that contained 100 μCi 5,6-^3^H-uracil, incubated at 30° for 10 mins and processed immediately for RNA. After electrophoresis, gels were soaked in Amplify Fluorographic Reagent (GE Healthcare) for 1 hour at room temperature, washed in water, dried, visualized by autoradiography to film at −80° and quantified as for Northern analysis.

### 2.3 Proteins

Wild-type and KE mutant strains were transformed with pNZ16 (pRS314 that carries *RPC128* with six N-terminal histidine residues followed by a FLAG tag, described in (Arimbasseri and Maraia, 2015). Chromosomal deletion of *RPC128* was achieved using *pFA6a-natNT2* and standard PCR-based methodology (Janke et al., 2004). Cells were batch grown in SC medium lacking leucine, uracil and tryptophan to OD 4.0 and Pol III was affinity-purified (Arimbasseri and Maraia, 2015). Pol III peaks were identified by Western blot for HA, pooled, dialyzed, concentrated and stored at −80°C. The Pol III preparations had comparable total protein yield and were normalized by the level of Flag-tagged C128 and C34 proteins detected by Western blotting.

The Tfc8TAP strain from the yeast TAP-tagged ORF library was engineered to overexpress Brf1 (approximately 50-fold) by substitution of the *natNT2 TEF1* promoter cassette (Janke et al., 2004) for the endogenous *BRF1* promoter. Whole cell extracts were prepared and chromatographed by size exclusion chromatography (Sephadex 200), cation exchange (Biorex 70, BRα step elution) and anion exchange (DEAE-Sephadex, step elution) to generate TFIIIB and TFIIIC fractions as described in (Nichols et al., 1990). The TFIIIC fraction was incubated in batch with 1 ml IgG Sepharose resin for 6 hours in 10 mM Tris-HCl pH 8, 150 mM NaCl, 10% glycerol, 1 ug/ml protease inhibitors (leupeptin, pepstatin A, aprotonin and 1mM PMSF). TFIIIC was recovered after overnight incubation with TEV protease (100 units, NEB). TFIIIC was empirically determined to be saturating and TFIIIB and Pol III activities were determined to be limiting by multiple-round transcription assays.

### 2.4 *In vitro* transcription

Factor-independent transcription on a tailed template was as described in (Arimbasseri and Maraia, 2015). Reactions contained EC buffer (40 mM Hepes-NaOH pH 8, 3 mM β-mercaptoethanol, 5% glycerol, 100 μg/ml BSA) with 7 mM MgCl_2_, 100 mM NaCl, 100 μM each ATP, CTP and UTP, 25 μM GTP and 10 μCi [α^32^P]GTP and 20 ng to 460 ng of template and were incubated for 30 mins at 22° in a total volume of 25μl. The 3’-tailed duplex template was prepared from gel-purified annealed oligos AGTO71 and AGTO72 as described in (Arimbasseri and Maraia, 2013). Reactions were started by addition of Pol III, stopped with 70 μl proteinase K solution (1 mg/ml in 10 mM Tris-HCl pH 7.5, 0.1% SDS), phenol/chloroform/isoamyl alcohol extracted and precipitated with 2 M ammonium acetate and ethanol (Willis et al., 1992). RNA was separated by 8% denaturing polyacrylamide gel electrophoresis, before exposure to phosphorimager as either a wet gel or after fixation and drying. Transcripts were quantified with ImageQuant software.

Non-specific transcription reactions were carried out for 30 mins at various temperatures in 50 μl with 1ug polydAdT copolymer, 1 mM ATP, 500 μM UTP and 0.5μCi [α^32^P]UTP in EC buffer plus 7 mM MgCl_2_ and 100mM NaCl (Huet et al., 1996). Reactions were started by addition of Pol III. Reactions were stopped and processed as described in (Roeder and Rutter, 1970) with 40μl 100 mM sodium pyrophosphate (pH 7) solution, containing 2 mg/ml salmon sperm DNA, 2 mg/ml BSA and 5 mM UTP, cooling followed by addition of 40 μl 5% SDS and 800 μl of cold 10% TCA, 40 mM sodium pyrophosphate. Acid-insoluble radioactivity was collected on Whatman GF/C filters, washed in 30 ml 5% TCA containing 40 mM sodium pyrophosphate, and counted in scintillation fluid.

Factor-dependent multiple-round transcription reactions were carried out for 60 min at 22° in 50µl with 20 mM Hepes-KOH (pH 8), 80 mM KCl, 2 mM DTT, 0.2 mM EDTA, 7 mM MgCl_2_, 10% glycerol and 100 to 250 ng plasmid DNA template. TFIIIB, TFIIIC and Pol III were added and the reaction initiated by addition of 1.2 mM ATP, 600 µM each CTP and UTP, 25 µM GTP with 10μCi [α^32^P]GTP. Reactions were stopped, processed and RNA separated and quantified as described for factor-independent transcription. Template DNAs were plasmids TZ1, which contains a G62C promoter up mutation in *SUP4* tRNA^Tyr(GUA)^ (Kassavetis et al., 1989), *SCR1*-C4T (Dieci et al., 2002), U6R (Whitehall et al., 1995), *sup9-e* (Willis et al., 1986), YEP13 containing *SUP53* tRNA^Leu(CAA)^ and *RPR1* in pRS423.

Single-round factor-dependent transcription reactions were assembled on TZ1 for 45 min at 22° with TFIIIB, TFIIIC, DNA and Pol III to form preinitiation complexes (Kassavetis et al., 1989). Stalled nascent 17 nucleotide transcripts were formed by the addition of 1.2 mM ATP, 600 µM CTP and 25 µM UTP and 10μCi [α^32^P]UTP for 5 mins. Time course experiments established that IIIB-IIIC-DNA-Pol III complex assembly was at equilibrium and that 17-mer synthesis was completed at or before 30 seconds for both wild-type and KE Pol III ternary complexes. Reactions were either stopped or extended to full-length transcripts by the addition of GTP and UTP to a final concentration of 600 µM with 300 µg/ml heparin (to prevent Pol III reinitiation (Kassavetis et al., 1989)) for an additional 5 to 10mins. Reactions were processed with 20 μg carrier tRNA as described and analyzed on 15% denaturing gels as above.

## 3. Results

### 3.1 Growth phenotypes of leukodystrophy mutations in the pore domain of Rpc160

POLR3A and its homologs are highly conserved across species and POLR3A HLD mutations identified to date are located throughout the entire protein (reviewed in (Wolf et al., 2014; Arimbasseri and Maraia, 2016b)). The most common disease-associated missense mutation in POLR3A, G672E, lies in a hotspot of missense residues that map to the pore domain of the enzyme; a region located below the floor of the DNA binding cleft that provides nucleotide access to the active site and where RNA is extruded during backtracking (Figure 1A) (Gnatt et al., 2001). We mapped pore domain POLR3A HLD mutations onto its *S. cerevisiae* homolog, Rpc160, in cryo-EM structures of the preinitiation complex and elongating Pol III (Figure 1A) (Hoffmann et al., 2015; Vorlander et al., 2018). The mutations are predicted to alter intramolecular interactions within Rpc160 and do not directly affect contacts with other Pol III subunits, subunits of TFIIIB or DNA (Figures 1A and 1B).

**Figure 1.**
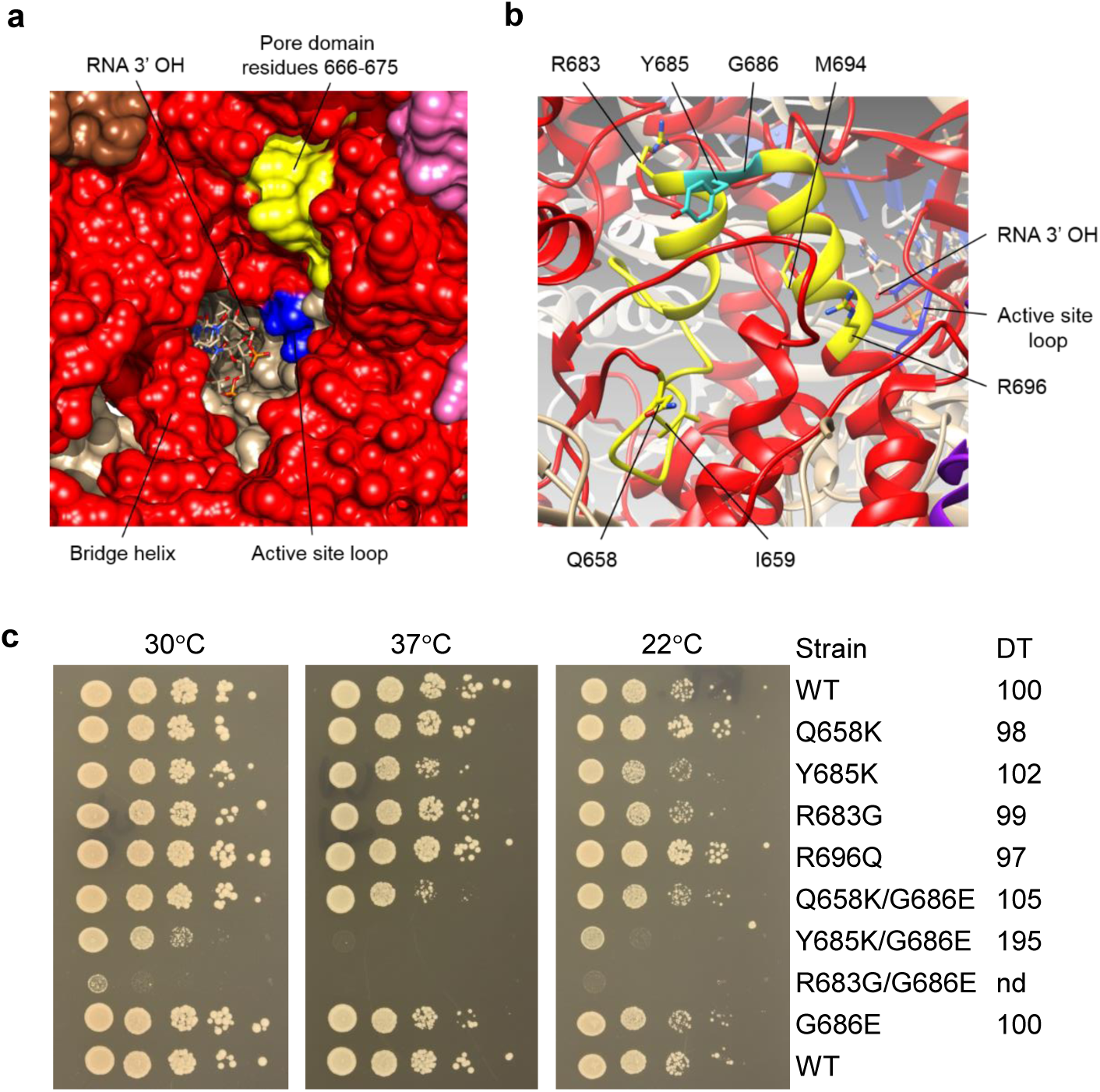
Rpc160 pore domain structures and mutant phenotypes. (a) Pol III elongating complex (PDB: 5FJ8), viewed from below the pore in the floor of the DNA binding cleft. Rpc160 (red) and its active site loop (residues 511-515, blue) are shown along with a small region of the Rpc160 pore domain (residues 666-675, yellow) that lies within a hotspot for HLD mutations (residues 658-696). Two helices within the hotspot (shown in panel b) lie behind residues 666-675. The bridge helix, RNA 3’ OH and parts of Rpb5 (brown) and Rpb8 (pink) are shown for orientation. (b) Seven disease-associated mutations in POLR3A are mapped onto the yeast Pol III elongating complex. Amino acids are numbered according to yeast Rpc160. The Y685K/G686E double mutant (abbreviated KE in the text, turquoise) maps to a tight turn between two α-helices (yellow). The active site loop of Rpc160 (blue) and RNA 3’ OH are shown in the background. (c) Growth phenotypes of wild-type, single and double mutant alleles of RPC160 in yeast. Ten-fold serial dilutions were spotted on SC-Leu plates and cells grown at 22°, 30° and 37°. The double mutant strain R696Q G686E was not viable. Doubling times (DT) in minutes at 30° in liquid media are indicated.

Five disease-associated missense mutations were engineered at the homologous positions in Rpc160 as single mutants, including the mutation G686E (corresponding to G672E in POLR3A). Each single mutant was also engineered as a double mutant in combination with G686E. The *RPC160* alleles were integrated at the *LEU2* locus in a *rpc160*Δ::*hphNT1* haploid strain where viability was maintained by a plasmid copy of the wild-type *RPC160* gene. The rescuing plasmid was evicted on FOA, and the mutant strains were evaluated for growth phenotypes and for Pol III transcription *in vivo*.

None of the single mutant strains exhibited a significant growth defect in liquid or solid media between 22° and 37° (Figure 1C). An analysis of total RNA showed that all of the single mutants had wild-type levels of mature tRNA and 5S rRNA (Figure S1A). The growth phenotypes of double mutant strains ranged from wild-type to lethal at 30°, with the three viable strains showing variable sensitivity to growth at 22° and 37°. The extremely slow-growing strain containing the R683G/G686E mutations exhibited a significant decrease in mature tRNA at the permissive temperature (Figure S1A). An intermediate growth defect was measured for the strain carrying the Y685K/G686E mutations. This double mutation, which we abbreviate to KE for simplicity, had a doubling time in liquid media almost twice that of the wild-type and its parental single mutant strains (Figure 1C) and was both cold-sensitive and heat-sensitive. The wild-type residues Y685 and G686 form a tight turn between two α-helices that pack onto other structural elements that position catalytically essential residues in the active site. We selected the KE mutant for further examination of Pol III transcription defects.

### 3.2 Compounding *in vivo* transcription defects in a *RPC160* double mutant

Northern analyses showed that the KE mutant had reduced steady state levels of a subset of Pol III transcripts *in vivo* compared to wild-type and its parental counterparts. The KE mutant showed a significant decrease in the steady state level of both precursor (pre-) and mature forms of *RPR1* RNA (the RNA component of nuclear RNase P, the enzyme that cleaves the 5’ leader sequence of pre-tRNAs) (Figure 2A). Pre- and mature levels of *SNR52* RNA (a non-essential snoRNA responsible for methylation of rRNA) were also decreased in the KE mutant (Figures 2B and 2C). In contrast, other Pol III-transcribed RNAs, *SCR1* (encoding the 7SL RNA component of the signal recognition particle required for co-translational insertion of proteins into the endoplasmic reticulum), *SNR6* (encoding the U6 snRNA component of the spliceosome) and *RDN5* (encoding 5S rRNA) showed no change in abundance (Figures 2E and 2F and Figure S1). The ratio of pre- to mature *RPR1* RNA was approximately one to one for all strains, suggesting that the low pre-*RPR1* RNA level in the KE mutant led to the reduced mature *RPR1* RNA level. Processing of the pre-*RPR1* RNA 5′-leader occurs after RNA assembly into the ribonucleoprotein complex (Srisawat et al., 2002) indicating that RNase P complexes might also be lower in the KE mutant. A probe specific to the 5’-end of pre-tRNA^Leu^ RNA showed full-length pre-tRNA and a very faint band corresponding to a spliced (but not 5’-end matured) intermediate in the wild-type and parental strains. The abundance of the partially processed tRNA intermediate was elevated in KE mutant RNA preparations and the level of full-length pre-tRNA and the intermediate varied significantly between biologically independent KE mutant RNA preparations (Figures 2D, 2G and S1), suggesting a stochastic reduction in RNase P activity in independent KE mutant cultures. The defect in pre-tRNA processing in the KE mutant was not sufficient to reduce the total mature tRNA level (Figure S1A).

As *RPR1* RNA is essential for viability in yeast (Lee et al., 1991b), we asked whether multiple gene copies could rescue the slow growth rate of the KE mutant. The doubling time of the KE mutant remained slow and its temperature sensitivity was unchanged by transformation with a multicopy *RPR1* plasmid (Figure 3A). In wild-type cells the increased *RPR1* gene copy number generated a five-fold increase in pre-*RPR1* RNA and a 50% increase in mature *RPR1* RNA. The effect on the KE mutant parental strains was less pronounced, generating a two to four-fold increase pre-*RPR1* RNA and a 20-25% increase in mature *RPR1* RNA. Increased *RPR1* gene copy number did not significantly alter pre- and mature *RPR1* RNA levels in the KE mutant and the activity of RNase P remained compromised (Figures S2 and 3B). The underlying basis for the absence of *RPR1* gene dosage effects on *RPR1* RNA in the KE mutant is unknown (see Discussion).

**Figure 2.**
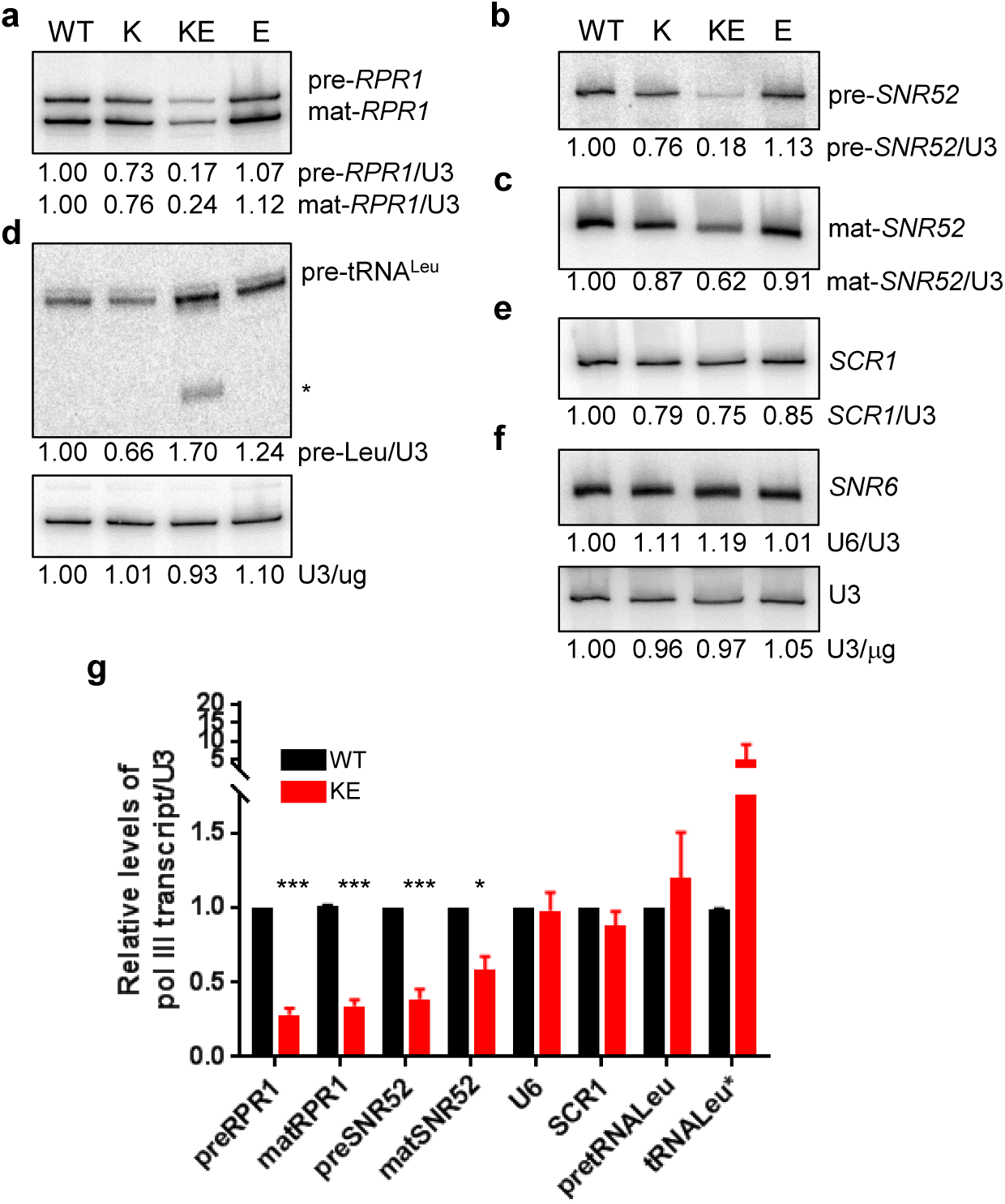
Northern analysis of Pol III transcripts. (a-f) Representative Northern blots of total RNA from wild-type, parental mutants with Y685K or G686E substitutions, and the double mutant KE (labelled WT, K, KE and E, respectively). Pre- and mature *RPR1* (a) and *SNR52* (b and c) RNAs, pre-tRNA^Leu^ (d), *SCR1* (e) and *SNR6* (f) RNA levels were quantified and normalized to the U3 snRNA loading controls. The amount of each RNA species is expressed relative to the wild-type value and indicated below each lane. Pre-tRNA^Leu^* is an intron-containing partially processed intermediate that retains its 5’leader. Data from multiple independent replicates are graphed in panel g. The error bars represent standard deviation (SD). p values are represented as *<0.05 and ***<0.0005.

**Figure 3.**
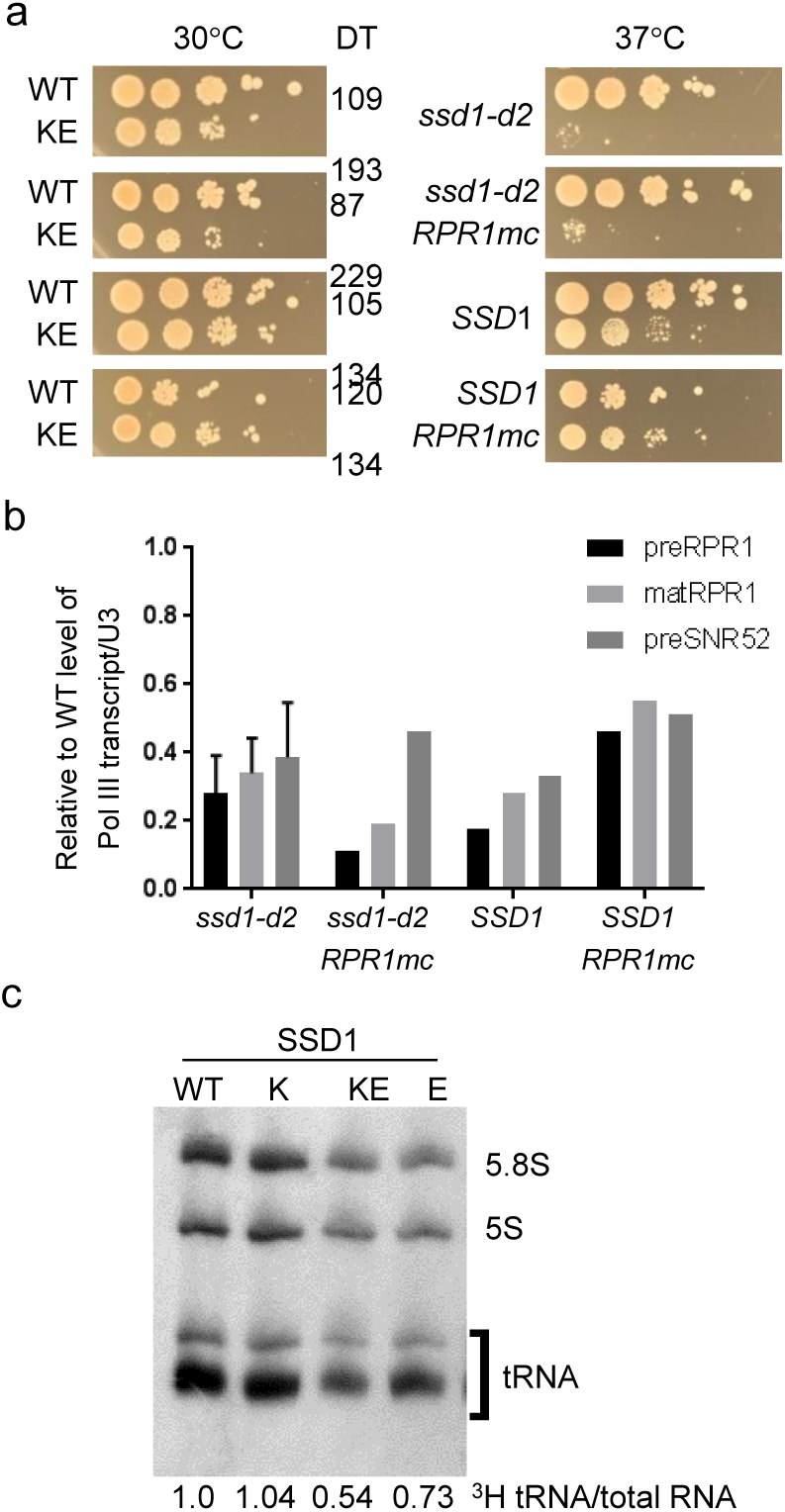
Growth and transcription defects of KE remain with wild-type *SSD1* and/or multicopy *RPR1*. (a) Growth phenotypes of wild-type and KE. Ten-fold serial dilutions were spotted on selective media. Plates were incubated at 30° and 37° for 3 days. The doubling times (DT) in minutes for each strain at 30° in selective liquid media determined by Bioscreen growth curves are indicated. (b) Graphical representation of Northern analyses of pre- and mature *RPR1* and *SNR52* transcripts in the KE mutant strains, normalized to U3 snRNA as a loading control. Levels of pre- and mature *RPR1* are expressed relative to the relevant wild-type strain. (c) ^3^H-uracil pulse-labelling of WT, YK, KE and GE strains containing wildtype *SSD1*. Total cpm incorporation/ug total RNA into each sample was comparable. Incorporation of ^3^Huracil into tRNA as a function of incorporation into total RNA is shown below each lane, expressed relative to wild-type.

Ectopic expression of the wild-type *SSD1* gene (*SSD1-V)* has long been known to suppress the conditional growth defects of mutations in Pols I, II and III in strain backgrounds (e.g. W303 used in this study) that contain the *ssd1-d2* allele, an early termination mutant of *SSD1* (Stettler *et al*., *1993*). Ssd1 is an RNA binding protein which affects the localization of specific mRNAs, limiting their translation (Hu *et al*., 2018) and Hsp104-mediated protein disaggregation (Mir *et al*., 2009). Introduction of the *SSD1-V* gene partially overcame the temperature-sensitive growth phenotype and suppressed the slow growth rate of the KE mutant at 30° (Figure 3A). However, *SSD1-V* did not reverse the defect in generating pre- and mature *RPR1* or pre-*SNR52* RNAs (Figures 3B and S2), nor did it alleviate the defect in RNase P processing of pre-tRNA^Leu^ (Figure S2). Incorporation of [^3^H]-uracil into total RNA in wild-type and KE mutant strains carrying *SSD1-V* was comparable, yet incorporation into mature tRNA by the KE mutant, calculated as either (i) the ratio of incorporation into 5.8S rRNA or (ii) the ratio of incorporation into total RNA, was low compared to wild-type values (Figure 3C). There was also a defect of [^3^H]-uracil incorporation into 5.8S rRNA relative to total RNA. Although the wild-type Ssd1 protein partially rescued the growth defects of the KE strain, the decrease in the abundance of pre- and mature *RPR1* and *SNR52* transcripts, and the reduction in neosynthesis of mature tRNAs, indicates that Ssd1 did not rescue the Pol III transcription defect.

### 3.3 Factor-dependent and factor-independent transcription is defective *in vitro*

Rpc160 protein abundance was equivalent in total cell extracts of both wild-type and KE mutant strains, whether extracts were prepared by native or denaturing methods (Figure S3A and B). This indicates that the half-life of Rpc160 is not significantly affected by the KE mutations, unlike several other mutations in Rpc160 which cause Pol III to be targeted by the sumoylation machinery and degraded (Wang et al., 2018).

Wild-type and KE strains were engineered to contain N-terminal His-tag and Flag epitopes on the second largest subunit of Pol III, Rpc128 (C128) (Arimbasseri and Maraia, 2015). Pol III was purified from both strains in comparable yield using immobilized metal affinity and anion exchange chromatography as per established protocols (Arimbasseri and Maraia, 2015). Affinity-purified Pol III was normalized for levels of C128 and Rpc34 (C34) proteins (Figure S3B) to quantify activity in factor-independent and factor-dependent *in vitro* transcription assays. Structures of Pol III show that C34, which is part of a trimeric C82/C34/C31 subcomplex, interacts with C160. Thus, the comparable recovery of C128 and C34 in the Pol III preparations reports the abundance of assembled Pol III complexes (Hoffmann et al., 2015; Abascal-Palacios et al., 2018; Han et al., 2018; Vorlander et al., 2018).

Factor-independent transcription of poly(dAdT), an alternating copolymer, was achieved with Pol III, ATP, and [α-^32^P]UTP. In this assay, KE Pol III was 55% and 40% as active as wild-type Pol III at 22° and 30°, respectively (Figure 4A). The activity of both polymerases was reduced at 37°, where transcription by KE Pol III was only 36% the level of wild-type Pol III. Factor-independent transcription was also assessed on a tailed oligonucleotide duplex template containing a 3’-single-stranded overhang and a 7bp T-run terminator sequence positioned 50 nucleotides from the transcription start (Arimbasseri and Maraia, 2013). KE Pol III was 56% to 59% as active as wild-type Pol III, at template concentrations ranging of 0.8 and 18 ng/μl (Figure 4B and data not shown). Both wild-type and KE Pol III exhibited a small but equivalent read-through of the terminator (22 ± 3% and 20 ± 3%, respectively, n = 8), which is attributed to dissociation of the C53/37/11 Pol III subcomplex that occurs during enzyme purification (Arimbasseri and Maraia, 2013).

**Figure 4.**
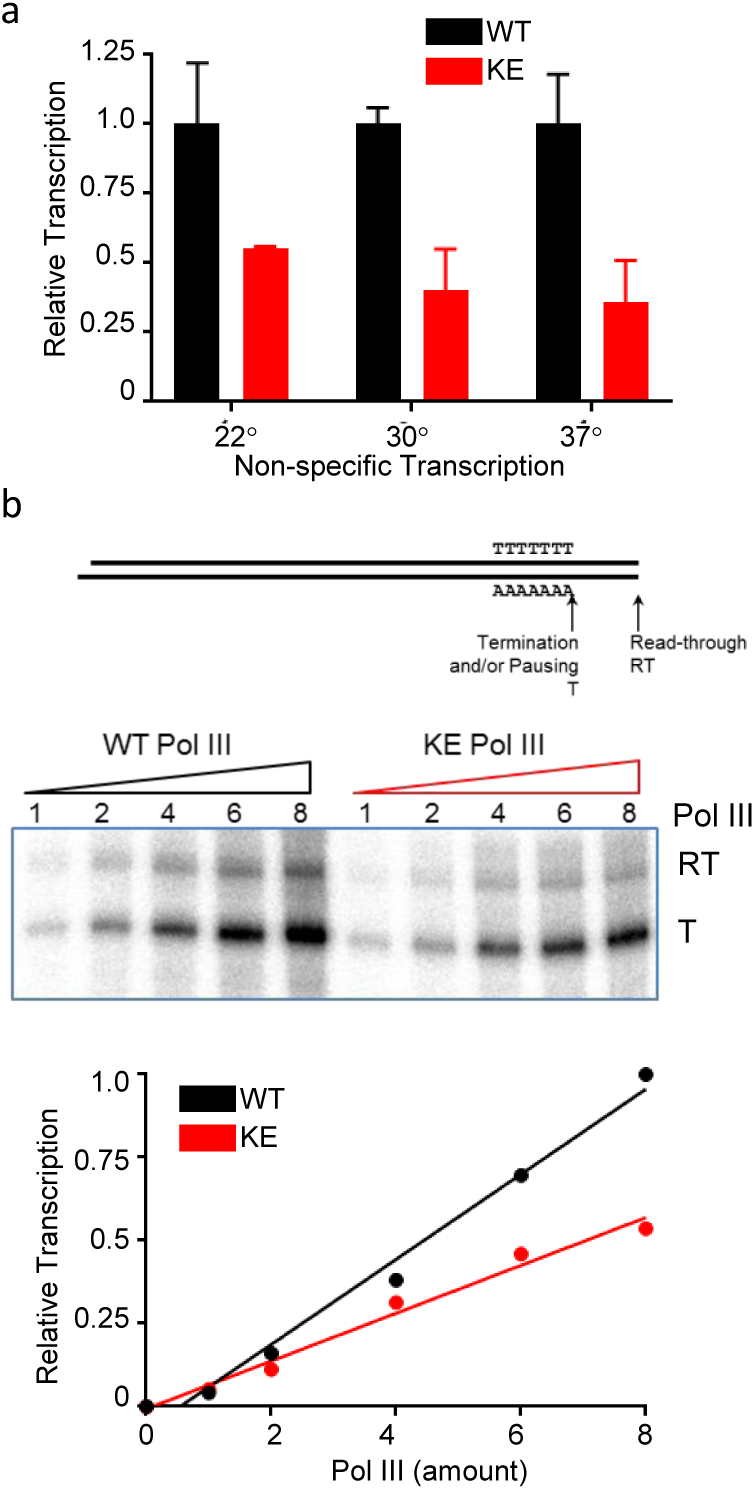
Non-specific transcription activity from wild-type and KE Pol III. (a) Non-specific transcription. Equal amounts of wild-type (black) and KE (red) affinity-purified Pol III was titrated into non-specific transcription reactions that used poly (dAdT) copolymer as a template. Transcription activity is expressed relative to that of wild-type Pol III at each temperature. KE Pol III was 55%, 40% and 36% as active as wild-type Pol III at 22, 30 and 37°, respectively. (b) Non-specific tailed template transcription. Equal amounts of wild-type (black) and KE (red) affinity-purified Pol III were titrated into non-specific transcription assays of a tailed oligo duplex. Transcription activity is expressed relative to that of wild-type Pol III. A representative titration at 250ng template is shown. KE Pol III had 59 to 56% the activity of wild-type Pol III on 20ng and 250ng template respectively. T, correctly terminated products; RT, read-through transcripts (22 ±3% and 20 ±3% read-through, for wildtype and KE).

Factor-dependent transcription by Pol III was assayed using a plasmid-borne *SUP4* tRNA^Tyr^ gene as a template. Affinity-purified yeast TFIIIC and a partially-purified yeast TFIIIB fraction were used to achieve high factor-dependent transcription activity. Both the DNA template and TFIIIC activity were in excess and Pol III was limiting in these assays. The activity of KE Pol III on *SUP4* DNA was 38 ± 10% (n = 9) the level of wild-type Pol III activity in multiple-round transcription assays (Figure 5A). A comparable defect was observed for transcription of a tRNA^Leu^ template (Figure 5B). Transcription was also compromised on Pol III-transcribed genes (*SCR1, RPR1* and *SNR6)* that deviate from the consensus sequence and/or spacing of type 2 promoter elements found in tRNA genes (Geiduschek and Tocchini-Valentini, 1988). Transcription of a dimeric *sup9-e* tRNA gene from *S*.*pombe*, which contains a tRNA^Ser^-tRNA_i_^Met^ gene where transcription is directed by the upstream tRNA (Willis et al., 1986), was also defective (Figure 5B). These results indicate that the activity of KE Pol III *in vitro* is broadly and equally defective on templates that vary in promoter strength and promoter element organization. Moreover, there were no shorter or longer transcripts generated on any template indicating normal start site selection and termination site recognition. The apparently normal steady-state level of mature tRNAs (Figure S1), *SCR1* and U6 RNAs (Figures 2E and F) in the KE mutant strain, coupled with defective Pol III activity *in vivo* and *in vitro*, indicates that there are homeostatic mechanisms at play to maintain a constant level of these transcripts *in vivo*.

**Figure 5.**
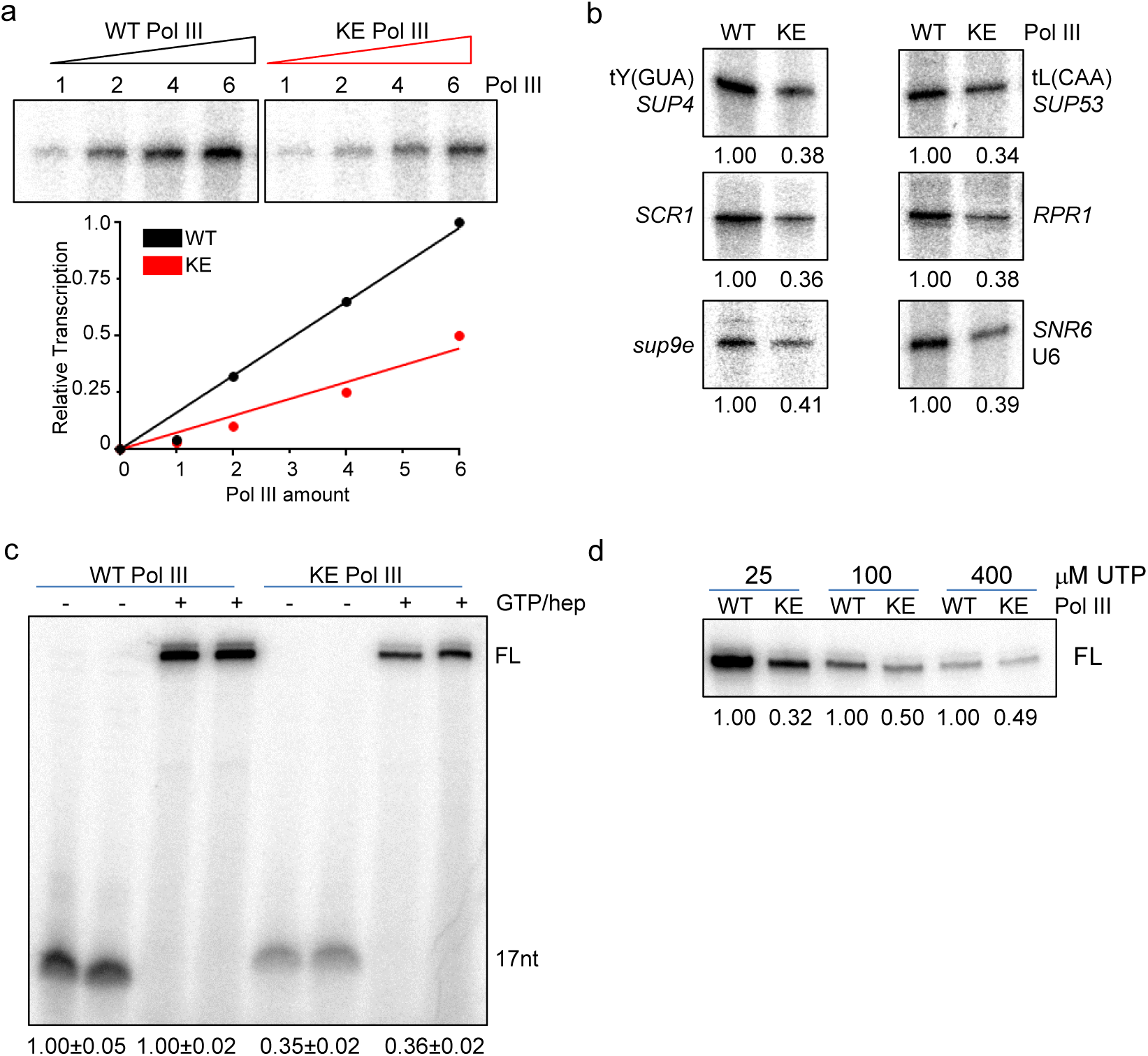
Factor-dependent transcription activity from wild-type and KE Pol III. (a) Multiple-round transcription of tRNA^Tyr^. Equal amounts of purified wild-type and KE Pol III were titrated into reconstituted transcription reactions that contained affinity-purified TFIIIC and yeast-fractionated TFIIIB. A representative multiple-round transcription image of SUP4 gene transcription is shown. KE Pol III had 38 ± 10% (n=9) the activity of wild-type Pol III. (b) Multiple-round transcription of other Pol III templates. tRNA^Leu^, *SCR1, RPR1, sup9e* and U6 genes were used to program transcription as in panel a, using fixed but equal amounts of each Pol III. The relative activity of KE Pol III is annotated under each template. (c) Single-round transcription of the SUP4 gene. A representative single-round transcription experiment is shown in which both the stalled nascent 17 nucleotide transcript (17-mer) and full-length transcript (FL) are resolved. Transcription was assessed by incorporation of ATP, CTP and [α-^32^P]UTP at 22° for 45mins generate (17-mers, lanes 1, 2, 5, 6). The addition of 600μM GTP, UTP and heparin for 5mins extended the labelled 17-mers to full-length transcripts and prevented additional rounds of transcription (lanes 3, 4, 7, 8). (d) Transcription with varying concentrations UTP. Full-length single-round transcription using 25, 100 and 400μM total UTP, with dilution of the [α-^32^P]UTP label. Transcription is expressed relative to the wild-type value set at 1.0 for each UTP concentration.

Factor-dependent single-round transcription assays were performed in which preinitiation complexes were assembled on *SUP4* DNA with TFIIIC, TFIIIB and Pol III. Addition of ATP, CTP and ^32^P-labelled UTP (25μM UTP, in the absence of GTP) allowed the formation of a ternary complex with a stalled nascent 17 nucleotide transcript (17-mer). 17-mer synthesis was lower for KE Pol III (averaging 35.2 ± 12% of the wildtype Pol III value, n = 9). Addition of unlabelled GTP and heparin to the stalled 17-mers allowed RNA synthesis to resume and limited Pol III elongation to a single-round of transcription (Kassavetis et al., 1989). These assays demonstrated that all labelled 17-mers formed by KE Pol III could be fully extended into full-length transcripts and that these generated a comparable differential; KE Pol III activity was 31 ± 6% the wild-type value (Figure 5C). No gross elongation or stalling phenotype was observed and KE Pol III transitioned from a stalled transcript into elongation mode to generate full-length RNA like wild-type (Figure 5C). The differential in single-round synthesis of full-length transcripts was maintained at high UTP concentrations (400μM) indicating that the defect in 17mer synthesis was not caused by hypersensitivity of KE Pol III to low UTP concentrations (Figure 5D). We noted that factor-dependent transcription by KE Pol III was modestly more defective than factor-independent transcription, suggesting that the TFIIIB, TFIIIC factors and/or supercoiled templates may impose further constraints on step(s) at which KE Pol III is defective. Together, the *in vitro* transcription experiments indicate that KE Pol III is globally defective at an early step in transcription on a variety of templates.

## 4. Discussion

The essential function of Pol III and the high conservation between POLR3A and Rpc160 (51% identical, 67% similar) suggested that pathological missense substitutions at orthologous positions in yeast would compromise Pol III transcription. This view was further supported by the fact that gene deletions involving the majority of Pol III subunits in yeast, all of which are essential, can be complemented by their human orthologs (Kachroo et al., 2015). HLD-associated missense mutations that map to the pore domain of Pol III were all able to complement the lethality of a yeast *RPC160* deletion. However, these single mutants exhibited no growth phenotypes and no measurable differences in the level of *RPR1* RNA or of a precursor tRNA that is normally sensitive to changes in Pol III transcription (Li et al., 2000b; Upadhya et al., 2002; Lee et al., 2015). The absence of transcription phenotypes for any single mutation in this region may result from yeast-specific amino acids within Rpc160 that buffer functional deficits in the protein and which negate, to a variable extent, the effect of a second mutation. Alternatively, as POLR3A is a substrate for both Hsp90 and co-chaperone binding in human cells (Taipale et al., 2014), species-specific differences in mutant protein folding, stability and complex assembly may also contribute to the suppression of a phenotype for the single mutations in yeast.

Purified KE Pol III has a global defect in function in both factor-independent and factor-dependent transcription *in vitro*. These findings and the location of the KE mutations on the opposite side of Pol III from its interface with TFIIIB (Abascal-Palacios et al., 2018; Vorlander et al., 2018), suggests that Pol III recruitment to the TFIIIB-DNA complex is unlikely to be affected by the mutations. Transcription by Pol III can be described as a series of fundamental steps including DNA binding to form a closed preinitation complex, DNA opening, transcription initiation, escape from abortive initiation, elongation, termination and facilitated recycling (reviewed in (Ramsay and Vannini, 2018)). Nucleotide incorporation directed by poly(dAdT) reports on all of these steps except terminator recognition and facilitated recycling. The tailed DNA template used in this work additionally reports on termination. Both assays indicated that KE Pol III was ∼60% as active as wild-type Pol III. KE Pol III was uniformly defective in factor-directed multiple-round transcription on a variety of Pol III-transcribed genes, with no indication of gene-specific defects. KE Pol III was also defective in the production of stalled nascent 17-mer transcripts, which encompass the DNA binding and opening, transcription initiation and escape from abortive initiation steps. KE Pol III was as competent as wild-type Pol III in supporting further polymerization of the 17-mer to produce a single full-length transcript. These *in vitro* transcription properties of KE Pol III are distinct from those of *rpc160-112* (a double substitution flanking the catalytic site) which showed a reduced elongation rate, increased pausing at intrinsic pause sites and slippage of nascent RNA in the stalled 17-mer (Dieci et al., 1995). As the activity of KE Pol III in single-round transcription and multiple-round transcription was comparably defective, at ∼40% of the wild-type activity, we suggest that KE Pol III may partition into two forms: one as competent for transcription as WT Pol III and the other form, functionally inactive. Whether the inactive KE Pol III population is defective in DNA binding or at an early step in transcription initiation has not been examined.

Although KE Pol III transcription *in vitro* was clearly defective, its defects *in vivo* were less apparent. The abundance of full-length pre-tRNA^Leu^ transcripts, an otherwise useful proxy for Pol III transcription, was not significantly altered in the KE mutant as tRNA processing was compromised. However, ^3^H-uracil incorporation into mature tRNAs was decreased compared to wild-type, clearly indicating a defect in global tRNA neosynthesis. Elevated nuclear exosome activity, estimated to specifically degrade pre-tRNA in wild-type cells such that less than 50% of pre-tRNA ends up as mature tRNA (Gudipati et al., 2012; Axhemi et al., 2020), could also contribute to the net decrease in labeled mature tRNA. However, as the level of total mature tRNA was not changed in the KE mutant relative to wild-type, the homeostatic mechanisms presumed to exist to maintain a constant level of mature tRNA in the cell are not affected in the KE mutant (Bonhoure et al., 2015). We have not assessed whether the composition of the tRNA pool, known to change in response to cell stress (Pang et al., 2014; Torrent et al., 2018), has been altered in the KE mutant.

The lower levels of pre- and mature *RPR1* and *SNR52* RNAs detected in the KE mutant *in vivo* likely stem from a combination of their non-canonical type 2 promoter structure, their unique processing and/or a requirement for specific RNA binding proteins to prevent their degradation (Lygerou et al., 1994; Srisawat et al., 2002; Preti et al., 2006; Palsule et al., 2019). Both *RPR1* and *SNR52* are synthesized as precursor RNAs from tRNA-like promoter elements that are positioned upstream of the mature RNAs and thus produce precursor RNAs with long 5’ leader sequences (Lee et al., 1991b; Harismendy et al., 2003; Lee et al., 2003; Preti et al., 2006). Although both genes contain A- and B-box internal promoter elements, the B-box in *RPR1* deviates from consensus (Lee et al., 1991a) and the distance between the boxes in *SNR52* is significantly longer than is typical. Additionally, the *SNR52* sequence contains a run of six T residues, normally a strong termination signal in the context of other Pol III-transcribed genes (Braglia et al., 2005; Arimbasseri and Maraia, 2016a). Transcription of *RPR1*, both *in vivo* and *in vitro*, is particularly sensitive to an internal deletion mutant of the Bdp1 component of TFIIIB (*bdp1Δ269-312*) (Ishiguro and Kassavetis, 2003) that is predicted to alter DNA binding by Bdp1 in the TFIIIB-Pol III preinitiation complex (Abascal-Palacios et al., 2018). Mature *RPR1* RNA (and RNase P function) remained low in the KE mutant despite additional copies of the *RPR1* gene, indicating elevated turnover of this pre-RNA. Cleavages to generate the mature 5’-ends of the *RPR1* and *SNR52* RNAs occur after assembly into their respective ribonucleoprotein complexes (Lygerou et al., 1994; Preti et al., 2006; Palsule et al., 2019), suggesting that low abundance of their specific binding proteins might underlie the low pre-RNA levels.

Pulse-labelling with ^3^H-uracil indicated that 5.8S rRNA biosynthesis was also decreased in the KE mutant. Crosstalk between defects in Pol III transcription and Pol I transcription is well documented. Mutations in *RPR1* RNA have been reported to affect 5.8S rRNA maturation *in vivo* (Chamberlain et al., 1996) and mutations in *RPC160* and 5S rRNA genes affect pre-rRNA processing (Hermann-Le Denmat et al., 1994; Dechampesme et al., 1999; Briand et al., 2001). It is unlikely that the reduction in mature *SNR52* RNA in the KE mutant significantly affects ribosome function, as a complete deletion of *SNR52* has limited phenotypic consequences (Esguerra et al., 2008). However, the reduction in 5.8S rRNA maturation and pre-rRNA processing may limit translation and underlie the slowed growth rate of the KE strain.

Although the *in vitro* transcription activity of KE Pol III was broadly defective, the growth phenotype of the KE strain was affected by the genetic background and the transcription defect of the KE mutation was detected at only a subset of Pol III-transcribed genes *in vivo*. In metazoans, the Pol III transcriptome varies in response to proliferation state, nutritional signaling, cell origin and stress (Dittmar et al., 2006; Coughlin et al., 2009; Canella et al., 2010; Canella et al., 2012; Ishimura et al., 2014; Carnevali et al., 2017; Mange et al., 2017; Yeganeh et al., 2019), and includes additional genes whose products affect pol II elongation (Peterlin et al., 2012), regulate autophagy (Horos et al., 2019) and, in humans, are implicated in translational repression and histone 3’-end maturation (Samson et al., 2018). The list of RNase P substrates is also expanded to include two long non-coding RNAs with pleiotropic activities (Wilusz, 2016). The additional complexity of the Pol III transcriptome, coupled with developmental, differentiation and tissue-specific gene expression, suggests that background modifiers of transcriptional defects will complicate readouts of defects in Pol III transcription in more complex organisms.

## 5. Conclusions

This work shows that the most frequent POLR3A HLD missense mutation, G672E, did not alter growth or Pol III transcription when engineered into its yeast homolog, Rpc160, as seen for the corresponding mutation in mice (Choquet et al., 2017). The addition of a second disease-associated mutation in the same region of Rpc160 caused intermediate defects in growth and Pol III transcription. This raises the possibility that introducing the corresponding double-mutant *Polr3a* gene in the mouse might generate a viable yet dysfunctional Pol III transcription phenotype and provide a mammalian model for the study of Pol III-associated leukodystrophy.

## Author Contributions

Each author has made substantial contributions to the work; has approved the submitted version; agrees to be personally accountable for the work. Methodology, C.L. and J.L; conceptualization, formal analysis, validation, investigation, resources, data curation, writing-review and editing, supervision, funding acquisition, R.D.M. and I.M.W; visualization, writing-original draft preparation, R.D.M; project administration, I.M.W.

## Funding

This work was supported by National Institutes of Health grants GM120358 to I.M.W. and HD097557 to R.D.M and I.M.W. and through an Albert Einstein Cancer Center core support grant, P30 CA013330.

## Acknowledgments

We thank Rich Maraia for the pNZ16 plasmid. We thank Emilio Merheb for his interest in this work and comments on the manuscript.

## Conflicts of Interest

The authors declare no conflict of interest.

## Supplementary Materials

The following are available:

Figure S1: Small RNA levels in hotspot single and double mutant strains.

Figure S2: Northern analysis of Pol III transcripts with strains containing *SSD1* and multicopy *RPR1*.

Figure S3: Pol III subunit association in KE extracts.

Table S1: Yeast strains.

Table S2: Oligo probe sequences.

## Notes

### Competing Interest Statement

The authors have declared no competing interest.

