## Supplemental Figs1_3 for "Functional characterization of *Polr3a* hypomyelinating leukodystrophy mutations in the *S. cerevisiae* homolog, *RPC160*"

**Supplementary Materials:**

**Table S1: Yeast strains.**

**Table S2: Oligo probe sequences.**

**Figure S1 Small RNA levels in hotspot single and double mutant strains.**

Five micrograms of total RNA was separated by denaturing polyacrylamide gel electrophoresis and stained with ethidium bromide to visualize 5.8SrRNA, 5SrRNA and tRNAs levels in wild-type, parental and double mutant strains. The loading amount of total RNA was empirically determined to be within the linear range for quantitation of all three RNA species. The tRNA and 5.8S rRNA signals were quantified by line-peak integration, the tRNA:5.8S rRNA ratio normalized to the wild-type value and indicated below each lane. Only the R683G/G686E mutant strain showed a significant decrease in mature tRNA levels. (b-d) Northern blots of total RNA from wild-type, single and double mutant strains. Pre- and mature *RPR1* (b) and pre-tRNA$^{Leu}$ RNAs are normalized to the U3 snRNA loading control (c). The amount of each RNA species is expressed relative to the wild-type value and indicated below each lane. Pre-tRNA$^{Leu}$* is an intron-containing partially processed intermediate that retains its 5'leader.

**Figure S2 Northern analysis of Pol III transcripts with strains containing *SSD1* and multicopy *RPR1*.**

Northern analyses of precursor and mature *RPR1* (a) and pretRNA$^{Leu}$ (b) transcripts in wild-type, KE and the parental mutant strains (labelled WT, K, KE and E) containing additional plasmids. W303 was grown with selection for an empty plasmid (labelled with *ssd1-d2*, the mutant *SSD1* allele in W303), a multi-copy plasmid containing the *RPR1* gene (labelled *RPR1*mc), a plasmid containing a wildtype *SSD1* gene (labelled *SSD1*) or both *RPR1* and *SSD1* plasmids (labelled *RPR1*mc, *SSD1*). Values are normalized to U3 snRNA as a loading control (c). Levels of precursor and mature *RPR1* RNA are expressed relative to the original wild-type strain.

**Figure S3 Pol III subunit association in KE extracts.**

(a) Western detection of Pol III in whole cell extracts from wild-type, single parental and KE strains. Extracts were prepared by TCA precipitation. The signal for C160 detected by its HA-tagged epitope and Rpl5 was detected with a polyclonal antibody (gift from John Warner) and ECL. C160 signal was normalized to that of Rpl5 and expressed relative to wild-type value. (b) Western detection of Pol III subunits in native extracts from wild-type and KE strains. Extracts from strains that contained Flag-tagged *RPC128* were prepared in isotonic buffer and detected with anti-Flag and HA antibodies. (c) Composition of affinity-purified Pol III. Equal volumes of WT and KE Pol III preparations were separated by SDS PAGE and western blots were sequentially detected with anti-HA and anti-Flag antibodies (6% gels) or C34 polyclonal antibody (10% gels).

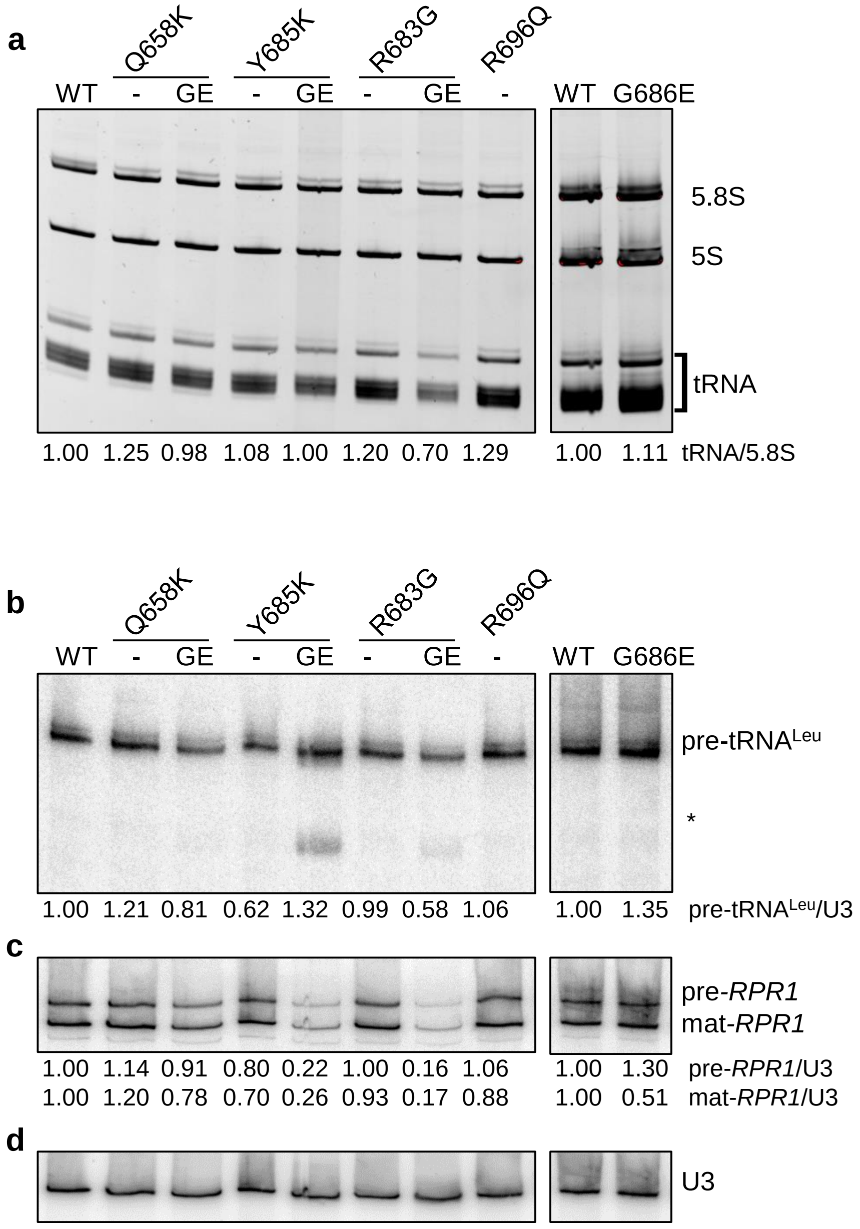
a
Q658K
Y685K
R683G
R696Q
WT - GE - GE - GE - WT G686E
5.8S
5S
tRNA
1.00 1.25 0.98 1.08 1.00 1.20 0.70 1.29 1.00 1.11 tRNA/5.8S
b
Q658K
Y685K
R683G
R696Q
WT - GE - GE - GE - WT G686E
pre-tRNALeu
*
1.00 1.21 0.81 0.62 1.32 0.99 0.58 1.06 1.00 1.35 pre-tRNALeu/U3
c
pre-RPR1
mat-RPR1
1.00 1.14 0.91 0.80 0.22 1.00 0.16 1.06 1.00 1.30 pre-RPR1/U3
1.00 1.20 0.78 0.70 0.26 0.93 0.17 0.88 1.00 0.51 mat-RPR1/U3
d
U3

Figure S1

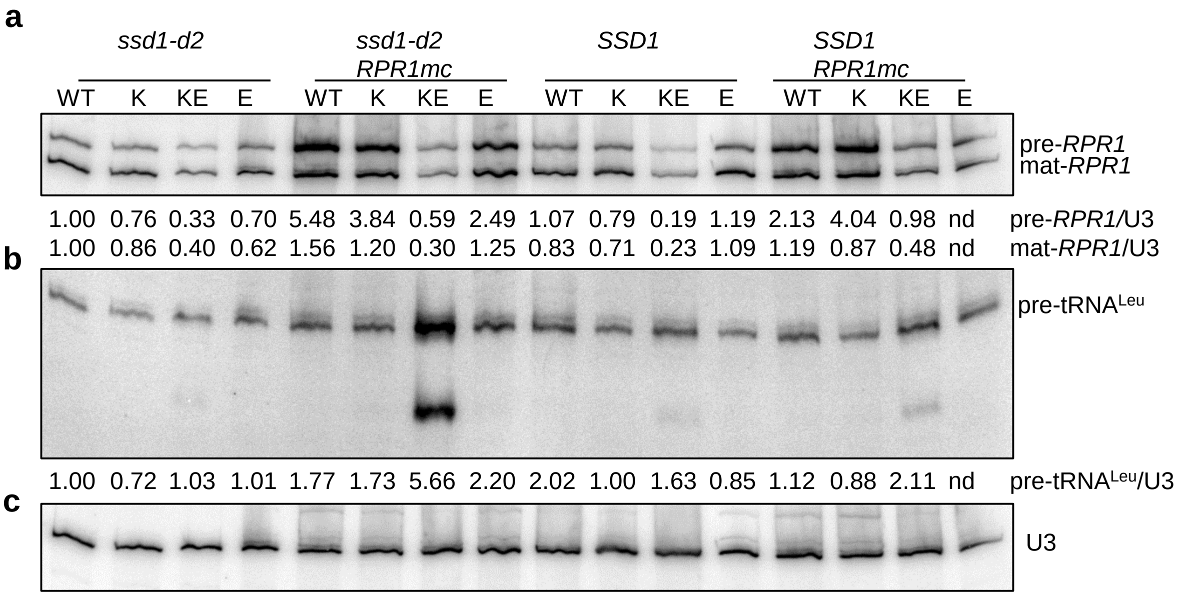
a
ssd1-d2
ssd1-d2 RPR1mc
SSD1
SSD1 RPR1mc
WT K KE E WT K KE E WT K KE E WT K KE E
pre-RPR1
mat-RPR1
1.00 0.76 0.33 0.70 5.48 3.84 0.59 2.49 1.07 0.79 0.19 1.19 2.13 4.04 0.98 nd pre-RPR1/U3
1.00 0.86 0.40 0.62 1.56 1.20 0.30 1.25 0.83 0.71 0.23 1.09 1.19 0.87 0.48 nd mat-RPR1/U3
b
pre-tRNALeu
1.00 0.72 1.03 1.01 1.77 1.73 5.66 2.20 2.02 1.00 1.63 0.85 1.12 0.88 2.11 nd pre-tRNALeu/U3
c
U3

Figure S2

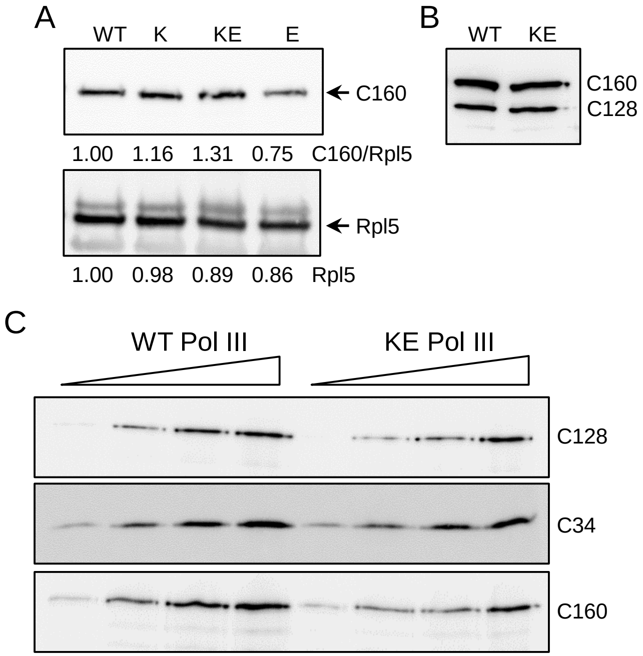
A
WT
K
KE
E
C160
1.00 1.16 1.31 0.75 C160/Rpl5
Rpl5
1.00 0.98 0.89 0.86 Rpl5
B
WT
KE
C160
C128
C
WT Pol III
KE Pol III
C128
C34
C160

Figure S3
